# VISIBLE: An imaging-driven system for sampling, biofabrication, and manipulation of complex biological models

**DOI:** 10.1101/2025.06.12.659321

**Authors:** Sudeep Joshi, Carmen Moreno-Gonzalez, Pacharaporn Suklai, Eugenia Carraro, Colin D.H. Ratcliffe, Giulia L.M. Boezio, George Konstantinou, Xavier Cano-Ferrer, Cathleen Hagemann, Thomas Kavanagh, Simon Ameer-beg, Albane Imbert, Erik Sahai, Francesco Saverio Tedesco, Andrea Serio

## Abstract

Complex in vitro models (CIVMs), including organoids, spheroids, and bioprinted constructs, have emerged as powerful platforms for recapitulating human tissue architecture and function. However, their inherent heterogeneity and dynamic nature pose significant challenges for standardization, reproducibility, and real-time manipulation. Here, we present VISIBLE (Versatile Imaging-Based Interactive Sampling and Bioprinting System), a modular, imaging-guided platform that integrates real-time monitoring with automated manipulation and 3D bioprinting that addresses these challenges. VISIBLE employs a dual-axis system comprising independently controlled tool-heads and microscope stages, enabling precise spatiotemporal interventions within live cultures. We demonstrate the system capabilities across a range of applications, including morphology- and function-based sampling of organoids and neurospheres, interactive 3D bioprinting, serial scratch assays, and real-time cell-cell interaction studies. Furthermore, we illustrate the system utility in translational contexts through the selective sampling and implantation of patient-derived xenograft organoids. VISIBLE supports long-term culture within an integrated incubation environment and accommodates interchangeable tool-heads for scalable, high-throughput workflows. By enabling closed-loop, feedback-controlled experimentation, VISIBLE addresses some of the critical limitations in existing CIVM platforms and offers a new versatile solution for a wide range of biomedical research applications. This work represents a significant step toward the development of autonomous experimental systems for complex tissue modelling and preclinical investigation.

## Introduction

Complex in vitro models (CIVMs)—encompassing organoids^1, 2^, spheroids^3^, micro-physiological “organ-on-chip” systems^4^, and bioprinted tissues^5, 6^ are engineered culture platforms designed to reproduce key structural and functional features of human organs more faithfully than conventional bi-dimensional (2D) monolayers. By incorporating multiple cell types within biomimetic extracellular matrices and, in many cases, integrating precise microfluidic control of nutrients, shear stress, and mechanical cues, CIVMs provide superior insight into human development, disease progression, and drug responses while decreasing reliance on animal models^7, 8^. Their increasing adoption across academic research, industry, and regulatory science reflects both a growing body of validation data and the pressing need for human-relevant test systems in bioscience and drug discovery. Over the last 10 years spheroids, organoids, organ-on-chip and other CIVMs have seen a significant increase in the number of studies that uses them as the main or one of the principal platforms for testing^9^. This upward trend underscored a global pivot toward complex human-relevant methodologies in bioscience, as a steady increase in adoption can be seen across industry^10^. This is also mirrored by changes in regulatory frameworks, which have recently started to acknowledge patient-derived organoids as acceptable non-clinical test systems^11^. CIVMs offer higher degrees of control for biological experiments, together with a human specific angle, and allow for real-time monitoring, manipulation of extracellular matrices, sampling of specific cell types, which are often unattainable in vivo. This ability to manipulate complex biology in a culture dish with ease has immensely benefitted our understanding of both fundamental biological questions and human disease, but it has also focused the technological development of tools and platforms in biosciences.

In fact, the rapid rise of complex human in vitro models, has been mirrored and enabled by equally swift advances in stem cell technologies^12^, bioprinting^13^, imaging^14, 15^, and laboratory automation^16^. These complex in vitro 3D models are often inherently heterogeneous, hence offers significant challenge towards standardization and reproducibility. Moreover, their generation and culture are complex and laborious, but in most cases also tied to the skills and capacity of the human operators. Recently developed systems to tackle the issue of reproducibility in complex in vitro experiments for instance, SpheroidPicker for 3D cell culture manipulation^17^ and imaging cell picker for morphology-based cell separation^18^ are limited in their operational abilities as they are developed for a singular functionality with almost no control and iterative manipulation abilities. 3D cell culturing via bioprinting ^19^ allow for the development of functional living tissues and organs, but lacks any manipulation ability during and after the printing process. Researchers are trying to push the boundaries further by integrating real-time imaging functionality termed as microscope-based 3D bioprinting^20^ for gaining visual control and modulation of the printing process. Although the modelling of these complex in vitro systems has become more automated, currently available systems still struggle to develop, control, and manipulate on-going cell culture experiments, especially in an iterative manner.

One of the main problems that persists in the technological platforms for CIVMs is the inherent variability in processing and lack of “on-the-fly” imaging-based intervention and manipulation capabilities. While current state-of-the-art systems, like fluorescent activated cell and organoid sorting (FACS)^21^ based on flow cytometry precisely isolate target cell populations from suspensions, they offer limited control and intervention, essentially functioning as a black box. Cell-positioning tools for microscopes are limited and present drawbacks in either throughput (e.g. micromanipulators^22^), set-up complexity (e.g. acoustic trapping^23^) or both (e.g. optical tweezers^24^). Recent advancements in microfluidic technology with integrated optics coupled with machine learning have enabled simultaneous deposition and monitoring of single-cell to gain insights into the mechanisms of cellular processes^25^ and to achieve higher efficiency in single-cell cloning^26^. The main limitation, however, is the lack of iterative intervention and manipulation capabilities. Continual usage of these systems seriously undermines further advancements of numerous in vitro biological experiments. To address these limitations, it is crucial to develop an advanced open-source system that allows higher-degree of control, real-time monitoring, and intervention for manipulation capabilities to leverage full potential of live complex in vitro experiments.

Here, we describe a platform that allows simultaneous handling, monitoring, and feedback-based intervention of complex in vitro experiments. We call this system Versatile Imaging-based Interactive Sampling and Bioprinting System (VISIBLE). This platform consists of a unique synergistic arrangement between two independently moving systems (top: a manipulation tool-head and bottom: a microscope stage) that allows for customization and thus facilitating greater control and manipulation capabilities. Continuous in-line imaging from the bottom microscope serves as feedback for the top manipulation tool head to allow for spatiotemporal intervention during live in vitro experiments. To validate the functional capabilities of our system, we demonstrate imaging-driven cell and organoid sampling (based on morphological differences and Ca^2+^ signalling), interactive 3D bioprinting (print-pause-analyse and place & pick), serial intervention (scratch assay and recovery), and real-time spatiotemporal interaction between muscle cells and neurons. Moreover, we were able to apply the VISIBLE’s capabilities to both in vitro, ex vivo, and in vivo preclinical settings.

## Results

### Engineering the VISIBLE system

The VISIBLE system comprises 3 modular units working in a synergetic arrangement for real-time monitoring, guided manipulation, and interaction with complex in vitro cell culture experiments. These are mirrored in the three main hardware components of the system: top section for deposition and manipulation, middle section for long-term cell culture and maintenance, and the bottom section for in-line imaging via an integrated microscope (**Fig.1A**). In the top section, the deposition and manipulation tool-head include two independent z-positioning linear actuators with 1 µm bidirectional precision, one dedicated for the deposition and the second dedicated for the visual feedback-based culture manipulation and sampling. The head consists of an in-house developed pneumatically-operated syringe extrusion system with pressure ranges regulated for single-cell sampling. The manipulation tool-heads house interchangeable syringes that support a range of nozzles (50-2000 µm) depending on the intended application. Like most conventional 3D bioprinting systems, our system uses 2 tool-heads, but we have additionally introduced an endoscopic camera firmly attached to them to map and precisely locate the sample position within the well-plate (**Fig.1B**). A macroscopic image of the sample from the endoscopic camera is used to position the microscopic stage, followed by a microscopic image of same the sample (**Fig.1B** centre and right panel). This novel feature of independently-controlled relative motion of microscope stage and manipulation tool-head movement allows for numerous versatile experiments.

**Figure 1:**
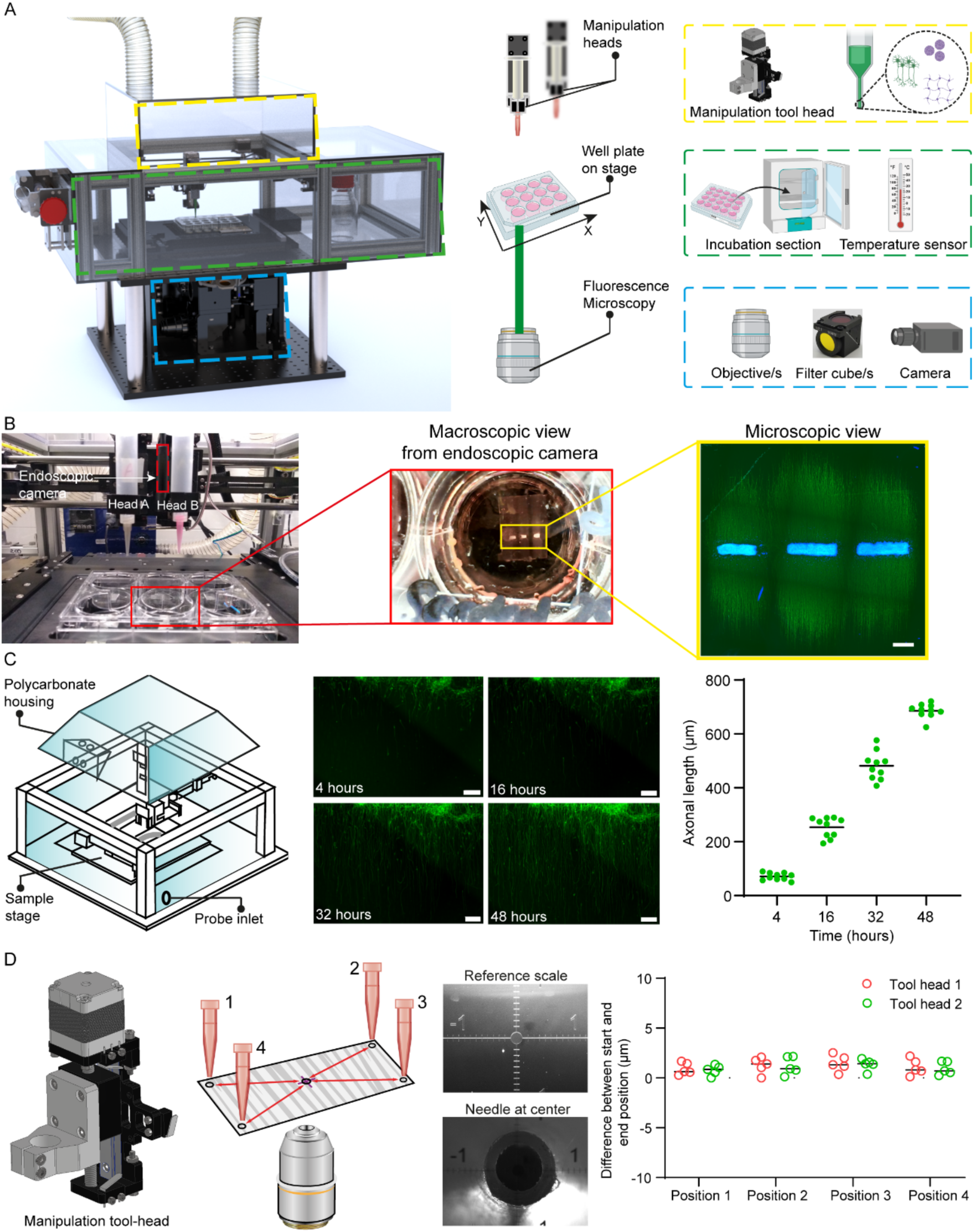
Engineering the VISIBLE system. (A) Schematic of the system demonstrating the imaging, incubation, and 3D biofabrication capabilities. (B) An image of a dual manipulation head (A & B) mounted with a syringe and a location detector endoscopic camera for identification of large area samples. Image showing a 6 well-plate placed on the controllable microscopic stage (inset), image of the sample captured from the top in-line endoscopic camera mounted on the tool-head for location detection of the sample under observation. An image of the same sample as captured from the inverted microscope of our system (Scale bar 1000 µm). (C) An image showing the incubation of the system and an associated motor neuron culture, wherein the extended processes are seen after 48 hours of incubation. A sequential image depicting the growth of processes from neurospheres for 48 hours to ensure suitability of the top-section as a cellular incubation chamber (Scale bar 200 µm). (D) Repeatability of the tool-head mounted on XY gantry. Bidirectional repeatability of the tool-head is in the range of ± 2 µm, hence ideal for single cell targeting and sampling with the VISIBLE system.

The entire top section consists mainly of manipulation tool-heads and an automated microscope stage, and it is encased in a polycarbonate enclosure (**Fig.1C**). This enclosure (volume of 0.12 m^3^) is designed to maintain an incubation environment (37 ℃ and 5% CO2) and also serves to maintain sterile conditions during manipulation. To test the long-term cellular incubation conditions within the system, we have conducted an experiment wherein human-derived induced pluripotent stem cells (hiPSCs) motor neurons are cultured for 48 hours. The growth dynamics of motor neurons were continuously monitored with the integrated imaging system (**Fig.1C**, centre and right panel). Cells were viable and extended processes (average length ∼720 µm) within the system’s incubation. This demonstrates the ability of our system to conduct long-term cell culture experiments with continuous monitoring of the events in vitro. The tool-head and its assembly on the XY gantry is designed to have sufficient precision to potentially conduct single-cell manipulation experiments. To test this, we performed a repeatability test and achieved a repeatable displacement of ± 2 µm (lesser than the neural cell diameter) from all the 4 corners of the test glass slide (**Fig.1D**). Within the enclosure, we also have a customized holder that serves as the repository for the samples collected by the manipulation tool-head during the experiments (supplementary **Fig.1E**).

### Precise live organoid sampling

Recent developments in the field of hiPSC differentiation, organoid technology and synthetic developmental biology have played a significant role in advancing our understanding of biological processes and disease states^27, 28^. While these techniques have been extensively optimised in recent years, a significant bottleneck for organoids-based in vitro studies remains: the inter- and intra-experimental organoid heterogeneity, for example in size and shape^29, 30^. Additionally, the existence of multiple differentiation protocols presents a significant challenge in replicating results across different researchers, laboratories, and even within the same lab^31^. Therefore, to efficiently characterise the variability in such dynamic cultures, continuous monitoring, and iterative intervention would be required to understand the cause behind the variation and ensure reproducibility. Currently, researchers use wide-diameter pipette tips to manually select and transfer the targeted organoids^32^ and fluorescent-based flow cytometry for sampling targeted populations in suspension^33^. However, these processes are labour-intensive and susceptible to human error, which can result in damaged cytoarchitecture and ultimately cell death. With these tools, continuous monitoring to track variability and targeted sampling of organoids for experimental fidelity remains a challenge.

VISIBLE allows the monitoring of growth dynamics, identification of morphological differences, and sampling of target organoids from heterogeneous populations for downstream analysis. Because of these capabilities, it represents an ideal system to address the challenges of variability and standardisation within organoids. To test this, the demonstrated the ability of our system to select and capture live organoids based on morphological features and fluorescent marker expression to decrease the inherent lack of reproducibility of organoid cultures. We first generated embryoid bodies (EBs) using custom-designed (400x400x200 µm) PDMS microwells^34^ and differentiated them into neuroectodermal organoids following a variation of a protocol described by Libby and colleagues^35^. In brief, hiPSC-derived EBs were treated with different concentrations of CHIR99021 (CHIR), a canonical Wnt activator, ranging from a low dosage of 0.8 µM to high dosage of 1.6 to 2.4 µM for the first two days to induce caudalization and thus a difference in symmetry breakage between the two populations^35^. After 48 hours, aggregoids were subjected to dual SMAD inhibition with SB431542 and LDN193189, to bias them towards the neuroectodermal lineage, while we continued CHIR treatment to activate Wnt signalling mimicking conditions in the posterior populations of the embryo (**Fig.2A**). As a result, we generated two different populations of organoids, with the high Wnt activation organoids showing distinctive symmetry breakage as expected^35^. To mimic an heterogenous culture with different morphologies, we mixed both low and high Wnt activation organoids and resuspended them in custom made PDMS sorting wells (**Fig.2A** and Supplementary Fig.2B). The PDMS sorting device was introduced into the VISIBLE system, wherein the system scanned the entire well and generated a high-resolution stitched image (**Fig.2B**). For this experiment, the organoids that broke symmetry were named as Population 2 and those that did not break symmetry were named as Population 1. Following the scan, the user can identify and select the organoids of interest and instruct the system to move to the precise location and lock the organoid using a nozzle. The VISIBLE system will then collect the organoid from the live suspension culture (Supplementary **video 1**) and transfer it to a desired tissue culture plates or Eppendorf tubes for downstream culturing or processing (Supplementary Fig.1E). This suction is based on the vacuum created by the hydraulic syringe pump attached to the tool head. The suction pressure is regulated in a manner that the pressure does not damage the organoid morphology. Previously reported studies on the lifting and transferring of organoids via Aspiration Assisted Bioprinting (AAB) saw a significant structural deformation of the lifted organoids. These macroscopic structural deformations result in significant alterations of the cellular organisation at microscopic levels^36^. Using VISIBLE, we were able to regulate this suction pressure via a precise custom-tailored syringe pump and collected the organoids without major alterations of cellular organisation at microscopic levels (Supplementary Fig.2D and **2E**). Next, the area of interest is again imaged to generate a high-resolution image to verify successful sampling of the targeted organoid without disturbing any nearby organoids (**Fig.2B**, rightmost panel). The VISIBLE system then performs several iterative steps to sample and collect all the organoids that have broken symmetry (Population 2) in a step-by-step process to appreciate the precise sampling capabilities of our system (Supplementary Fig.2C).

**Figure 2:**
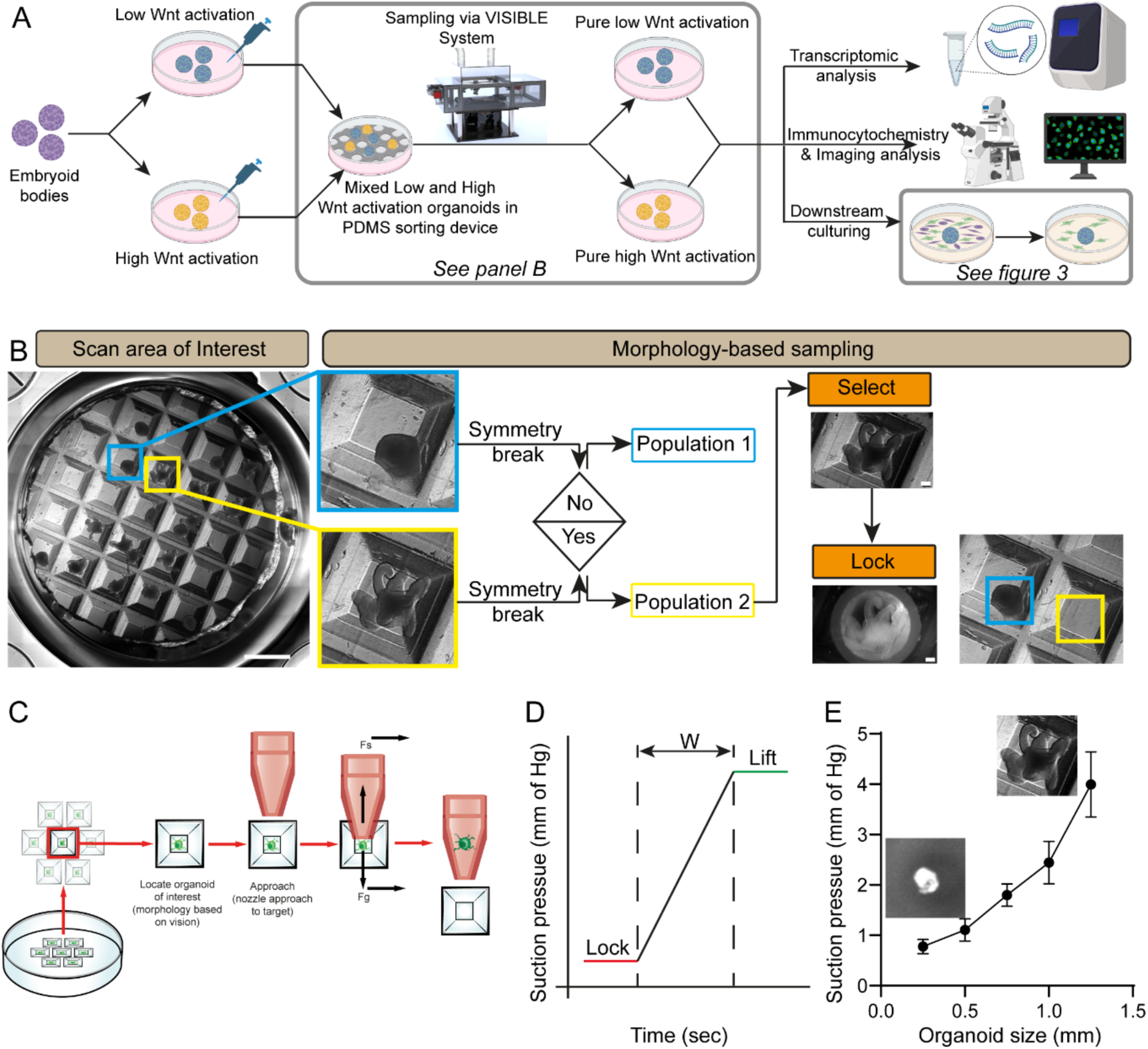
Morphology-based sampling of suspended neuroectodermal organoids. (A) Schematic of the experimental design wherein neuroectodermal organoids are treated with different concentrations of CHIR molecule resulting in low and high Wnt activation populations. These are mixed in equal proportions to generate a heterogeneous culture to be sampled by the VISIBLE system followed by downstream analysis of the collected organoids. (B) Process flow chart of the operation of the VISIBLE system wherein the system scans the entire area of interest: a PDMS macrowell containing a mixed organoid population in suspension (Scale bar 1 mm). Identification of organoids that break symmetry as “Population 2” and those that do not as “Population 1” for the collection for further analysis. The action sequence of the VISIBLE system, wherein the system selects the organoid of interest and a nozzle locks the organoid for collection. This is followed by another scan of the area of interest to confirm the organoid collection. (C) A schematic showing the procedural steps for locating, moving, and locking up the organoid of interest by the VISIBLE system. (D) A graph showing the relation between suction pressure and time, wherein W varies depending on the dimension and mass of the organoid. (E) Variation of suction pressure versus organoid size indicates a linear dependence.

To characterise their viability, preserved structure and test the feasibility of transferring organoids without damage, these sampled organoid populations were then stained with DAPI, showing well-preserved organoid cytoarchitecture of sampled Population 1 and 2 (**Fig.3A**). Likewise, population 1 was also collected and stained, revealing similar results. In parallel, we extracted RNA from a proportion of the sampled organoids and original cultures to perform gene expression analysis. Relative gene expression shows that the VISIBLE-picked high Wnt population presents similar levels of neuroectodermal (*SOX1, SOX2, PAX6*), posterior identity (*HOXC9*) and somitic and presomitic markers (*TBXT, TBX6*) as the pure non-sorted high Wnt population, suggesting that the system successfully sampled only the desired organoids and preserved them for successful RNA collection. Moreover, a subset of the sampled organoids was also picked and plated in a tissue culture well to assess the viability of transferred organoids. After 2 days of culturing, the plated organoids showed migratory cell populations (**Fig.3B**), suggesting that the organoids survived the transfer process and did not show any signs of contamination. Taken together, these experiments serve as a proof-of-principle that the VISIBLE system is able to select, pick, and collect complex 3D human organoids while preserving their structural and cellular integrity.

**Figure 3:**
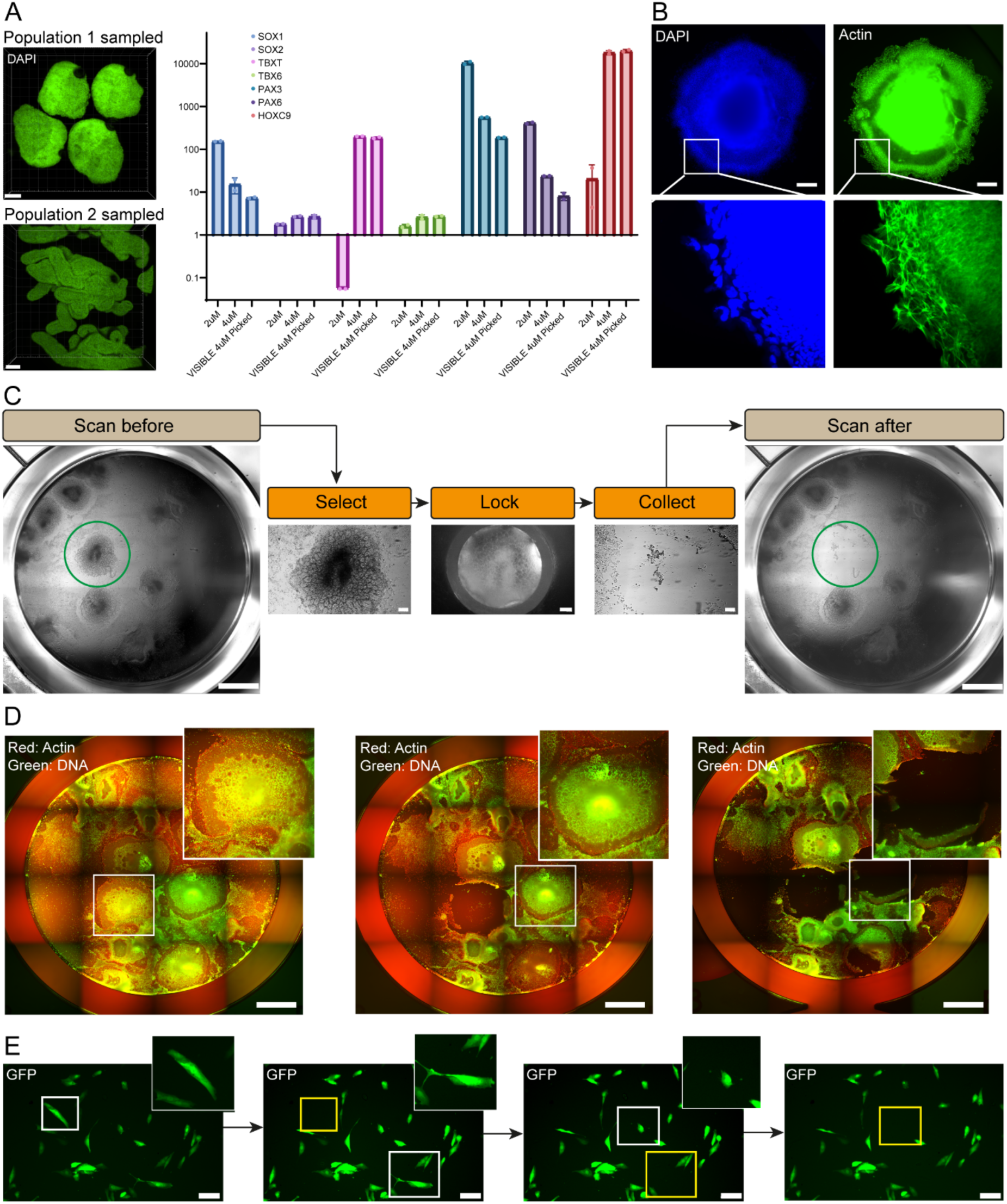
Morphology-based sampling of adherent organoids. (A) ICC studies of population 1 and population 2 organoids sampled via the VISIBLE system and a graph showing relative gene expressions (Scale bar 2µm). (B) An organoid plated via VISIBLE system remains viable and after 2 days in culture, the plated organoids showed migratory cell populations. (Scale bar 1µm). (C) The system also has the capability for sampling of adherent organoids and their migratory populations with high precision managing to not disturb nearby organoids (Scale bar 1 mm). Here, Actin filaments were spotted via a microscopic image and the attached organoids were collected via executing a similar process chain. (D) VISIBLE system also allows for the use of multiple fluorophores to detect numerous proteins of interest and sampling can be directed based upon protein presence (Scale bar 1mm). (E) Iterative sampling of adherent GFP+ skeletal muscle cells (∼50 µm) to demonstrate the VISIBLE system precise collection capability (Scale bar 100 µm).

### Sampling of organoids in adherent cultures

Next, we tested its capability for suction-based collection in adherent cultures as the magnitude of suction pressure differs because organoids are attached to the plate. Following the previous experiment, we plated a mixed population of “low” and “high” Wnt activation organoids into a tissue culture plate and cultured them for an additional 6 days. Due to their intrinsic differences created during the initial induction phase, these organoids exhibited clear morphological differences when attached and generated distinct migratory cell populations in their vicinity (**Fig.3C**). These differences in the growth pattern were of interest in this round of experiments and individual organoids were selected by the VISIBLE system for further sampling. Our system enables the controlled lifting, transfer, and replating of organoids. Success of this transfer is confirmed by the preserved cytoarchitecture of the targeted organoid and its viability after 48 hours of transfer. Different concentration of CHIR resulted in different sized organoids and mass^36^. As a result, the suction pressure required for lifting of organoid was directly proportional to the size of organoid (**Fig.2E**), similar observation is also corroborated by other research studies^36, 37^.

One of the main aims in designing the VISIBLE system was to enable imaging-driven manipulation, granted by the coupled and co-registered microscopy and gantry systems within the instrument. This translates into the ability to visualise a range of fluorophores that label specific cellular components, and use them as parameters in the target identification for sampling based on features of interest, for example here with fluorescently lineage-labelled organoids (**Fig.3D** and supplementary **video 2**). This particular imaging feature followed by function-based sampling of adherent organoids is not possible with the current systems such as cell-sorting devices and organoid sorters. Furthermore, to demonstrate the precision of hydraulic suction-based sampling via the VISIBLE system, we have collected single skeletal muscle cells (∼40-80 µm) from a dense culture, iteratively (**Fig.3E**).

### Function-based sampling

Another key factor that presents a high degree of variability within a complex cultures is the functionality of different cells. We wanted to test the ability of VISIBLE to target and sample based on functional profiles. Neurons were differentiated via doxycycline inducible expression of the NGN2 gene integrated into an hiPSC line (KOLF2.1J) using the piggyBac transposon system. (**Fig.4A**). Neurospheres were subsequently generated as previously described^34^ with modifications. These neurospheres were plated on a Matrigel-coated culture plate and maintained for 16 days to allow for maturation. Maturing glutamatergic neurons especially in a complex sphere, connect to each other and exhibit calcium spikes and transients, but each neurosphere will have a different degree of this connectivity resulting in altered calcium activity and different maturity levels across the same cultures. We used a calcium dye to assess these functional maturation differences and actively select single neurospheres to isolate. We chose this experimental set up as it is widely applicable to different CIVMs cultures and experimental pipelines. As a first step, the system generates a stitched image of the entire well to locate the different neurospheres and log their position, and subsequently higher resolution videos can be captured to assess their level of activity based on the Ca^2+^ fluctuations. For this experiment, we selected 6 randomly placed neurospheres from the generated stitched image (**Fig.4B (I)**). Multiple positions within a selected neurosphere were identified, and neuronal activity was recorded over a period of 5 minutes with 1 second intervals, using 488 nm excitation (Supplementary Fig.3B and **C**). The neurospheres that showed higher numbers of Ca^2+^ peaks indicate higher neuronal activity and were selected for sampling using the VISIBLE system (**Fig.4B (II)**). Once the neurospheres of interest were selected, the system precisely moved the tool-head fitted with an appropriate nozzle diameter (520 µm) via Z-axis movement to lock the target (**Fig.4B (III)**), which is then collected via a controlled suction mechanism. The images illustrate the precision of VISIBLE system in sampling a 500 µm neurosphere, wherein even the processes of nearby neurospheres stayed intact (**Fig.4B (III)**). Such a precise imaging-guided targeting of neurospheres based on maturation and functional parameters within a single instrument with real-time feedback is highly desirable and represents an enabling technology for more complex and heterogeneous experiments, which has not been reported by any other system in literature. We were able to perform the process of neuronal activity-based sampling in an iterative way and repeat it for the remaining 3 active neurospheres, followed by imaging of the area of interest again for confirmation. The collected samples can then be used for downstream applications such as imaging and transcriptomics as previously demonstrated during the previous organoid sampling experiments. Overall, the VISIBLE system was able to perform function-based sampling in live cultures, detecting and sampling neurospheres with higher Ca^2+^ dynamics: a hallmark of their maturation. This demonstrates our system capability to handle imaging-based sampling based on functional profiles in complex adherent cultures, which is a unique feature that can be applied across several experimental paradigms.

**Figure 4:**
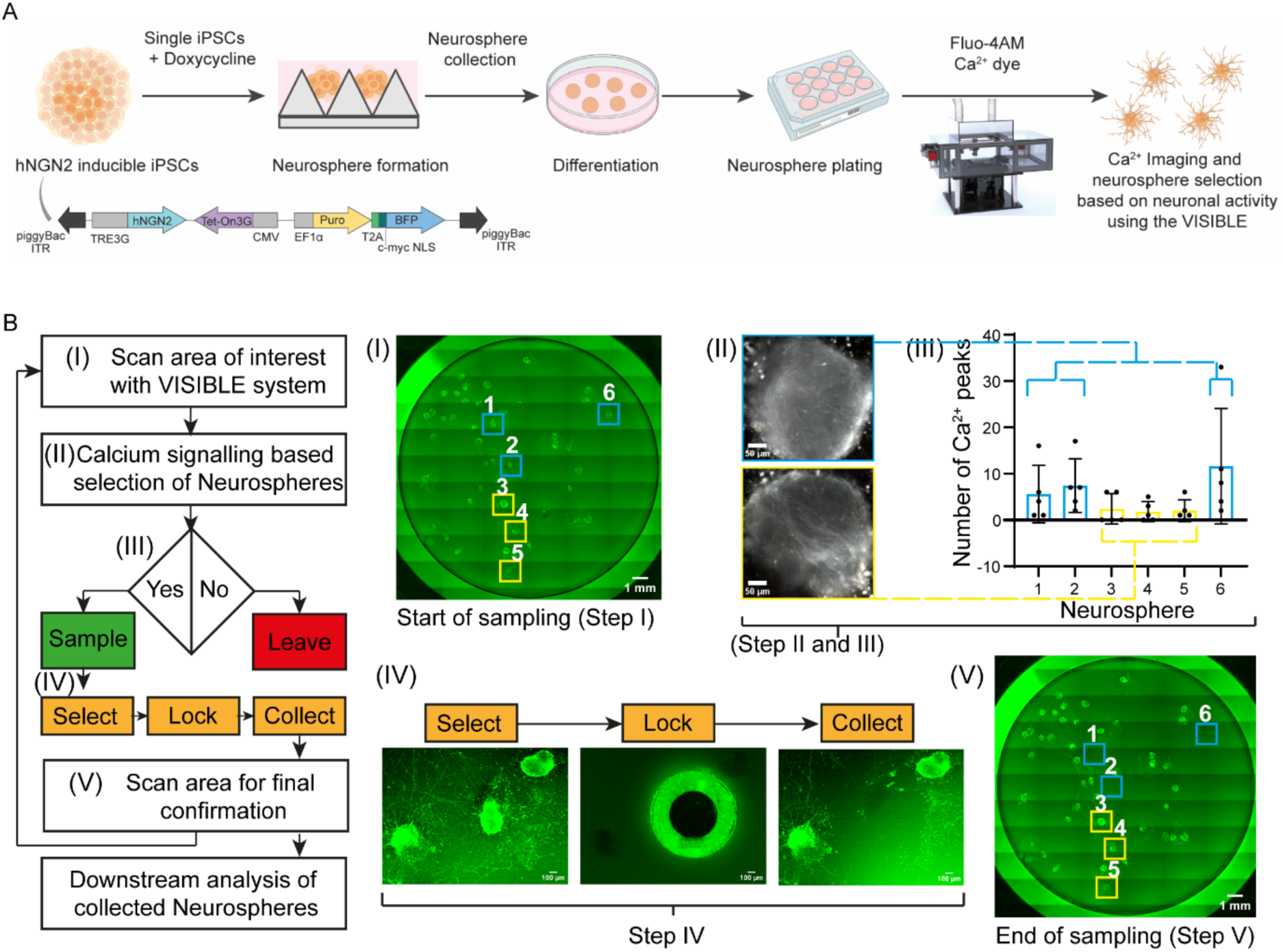
Function-based sampling of adherent neurospheres in complex culture. (A) Schematic of hiPSC derived neurosphere formation and plating in a 12-well plate for interactive calcium signalling experiment with the VISIBLE system. (B) Process flow chart of the operation of VISIBLE wherein (I) The system scans the entire area of interest with a number of neurospheres attached to Matrigel-coated plate (Scale bar 1 mm). (II) Identification of neurospheres that exhibit calcium signals and whose number of peaks are more than 5 from 10 ROIs. (III) The VISIBLE system selects the neurosphere of interest (exhibiting more than 5 peaks) and a nozzle locks it for collection. (IV) The system scans the well again to confirm that the neurosphere of interest is collected. The collected neurosphere is plated into another well for further downstream analysis. (V) Final scan of the well showing the selected neurospheres with higher numbers of calcium peaks (1,2, and 6), which were collected sampled out and the ones with lesser peaks (3,4, and 5) remain intact (Scale bar 1mm).

### Interactive 3D bioprinting and serial intervention for spatiotemporal experiments

Another major obstacle to further advance in vitro cultures is the interaction with live cultures based on real-time imaging feedback. For example, studying complex systems such as neural circuits, developmental niches, and tumour microenvironments ideally requires a platform capable of building intricate architectures of live cells and extracellular matrix (ECM), while also enabling dynamic manipulation of the culture’s behaviour and composition. Moreover, real world screening or phenotyping experimental pipeline would ideally require to interact with the same samples in an iterative manner, for example to sample multiple timepoints across one well or deposit different cell populations at different times. Currently available 3D bioprinters can print 3D structures ranging from mm to cm^5, 6^ and generally do not have any real-time feedback capability, as they perform commands based on the design file provided by the user without any possibility of intervention and manipulation. This is problematic for complex biological cultures, which are by definition very dynamic in nature. With our system’s real-time imaging-driven features, we have greater control as the system allows for feedback-based iterative printing and analysis ability within live experiments.

The VISIBLE’s XY tool-head gantry and microscope stage movements are independent of each other and co-register on the sample. This unique feature allows to generate real-time images and adapt the tool-head actions so that it can iteratively interact with the complex culture based on the imaging feedback. To demonstrate this, we performed some proof-of-principle bioprinting tests. We designed a printing experiment based on bioprinting iPSC-derived cell population within alginate gels (see M&Ms) in a predefined architecture. We split the printing of the desired geometry into 4 different segments, and showed we were able to print separate parts of it, then image the resulted cell-embedded hydrogel constructs, to check for any abnormalities (**Fig.5A** and Supplementary **video 3**) before printing the following sections of the designs. The imaging-driven feature of VISIBLE allowed us to iteratively print and analyse each step of the 3D hydrogel mesh, which would allow to adapt the printing protocol “on the fly” to reduce overall failures. We then tested the iterative printing and sampling of the printed construct with the VISIBLE system. For this we generated a sequential dot printing of hydrogel matrix, and afterwards selected regions to be sampled without disturbing the nearby dots (**Fig.5B**). This showed that the pneumatic sampling mechanism of VISIBLE can be combined with the bioprinting and is capable of complex interactions with 3D cultures. Moreover, we were able to test the viability of cells printed in hydrogel with the VISIBLE system (**Fig.5C**) showing that the printed constructs are viable 2 days after construction in vitro. This also demonstrates that the hydraulic printing mechanism exerts appropriate amount of pressure for hydrogel printing without damaging the cells in the printing process.

**Figure 5:**
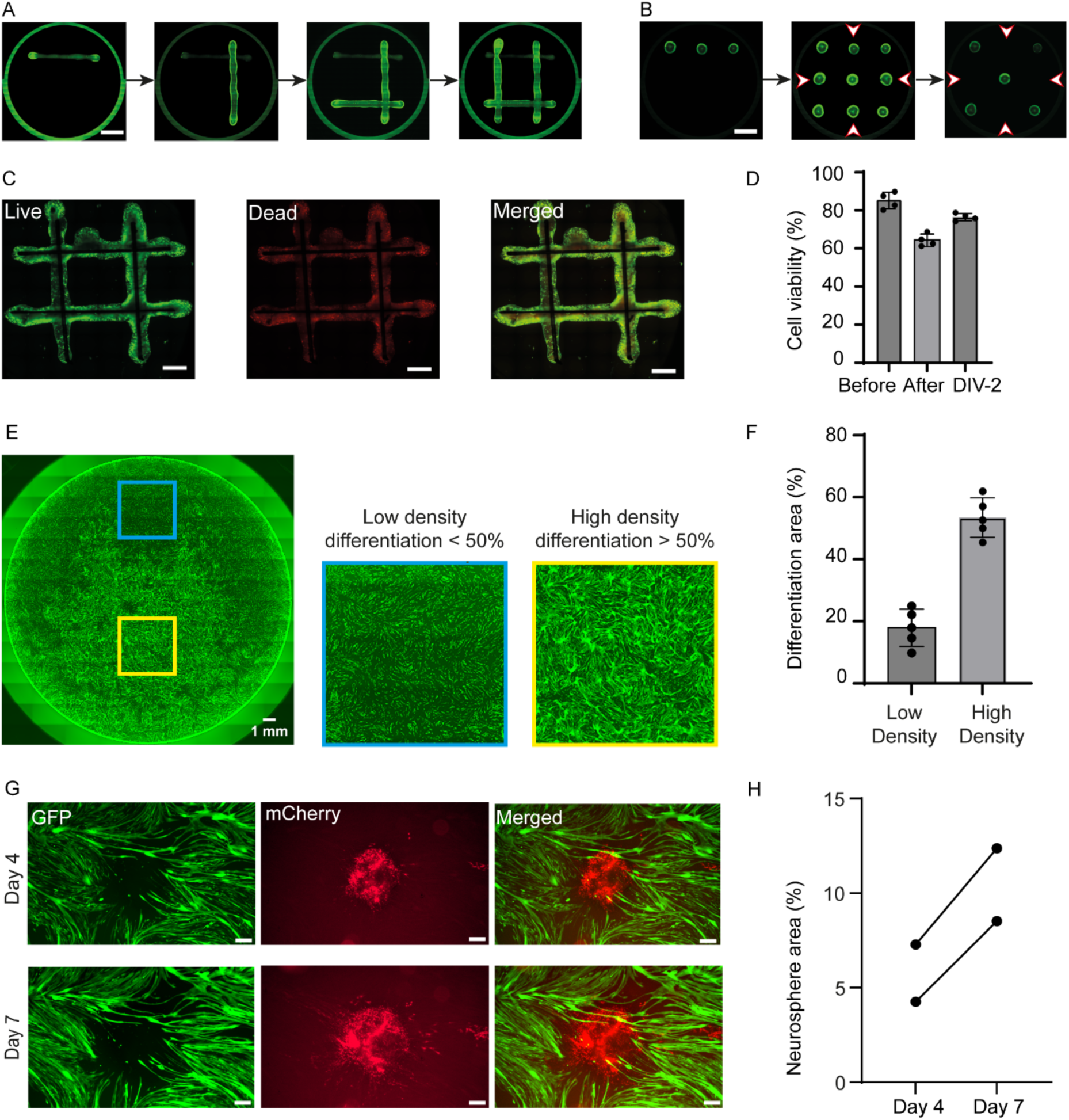
Interactive 3D bioprinting and serial intervention for spatiotemporal experiments. (A) Serial intervention 3D printing of hydrogel, wherein print-pause-analyse-print procedure is demonstrated, allowing for live interaction based upon a previous print (scale bar 1 mm). (B) A 3D printing of dot-patterned hydrogel, followed by selection and collection of the region of interest (scale bar 1 mm). (C and D) 3D bioprinting of live cells within the alginate-based hydrogel matrix showcasing viability after the printing process to establish that the process of hydraulic-based extrusion of cell-embedded hydrogel is biocompatible (scale bar 1 mm). (E and F) Scan of a well of a 12 well plate cultured with differentiated muscle cells followed by the selection of 2 areas with different differentiation densities of muscle cells and respective quantification (G and H) Co-culturing of neurospheres deposited on high-density area of muscle cells for 1 week, demonstrating interaction between processes of motor neurons with the differentiated muscle cells and increased coverage area (Scale bar 200 µm).

The imaging-based real-time feedback capabilities of VISIBLE allows for dynamic responses in creating and manipulating complex cultures. This becomes very important and useful in the context of constructing complex cultures with multiple cell types differentiating at different speeds which are needed to form a functional unit. An example of this is co-cultures of iPSC-derived motor neurons (MNs) and skeletal muscle cells, for the purpose of creating neuromuscular junctions: only highly mature muscle fibres would successfully interface with MNs. However, within one culture well there can be high-levels of variability of differentiation efficiency and myofibers maturation, so the user would need to select where to deposit MNs directly. This paradigm is also transposable to any other complex coculture with multiple cell types or dynamic cell states. To demonstrate the ability of our system to tackle this, we have grown human myoblasts on a Matrigel-coated tissue culture plate and observed the differentiation pattern over a week of culture within the VISIBLE system. Myoblast fusion and differentiation into multinucleated myotubes is triggered by cell-cell contact; as a result, heterogeneity in cell culture conditions (e.g., density) results into areas of the culture dish with distinct differentiation efficiency. VISIBLE allowed us to scan the entire well to identify the areas with relatively higher differentiation (areas with differentiation >50%) in real time based on the morphology and size of the myotubes (**Fig.5E** and **5F**). These highly-differentiated areas were selected to then deposit iPSC-derived neurospheres containing motor neurons. Neurospheres embedded in hydrogel were deposited on the selected locations and allowed to interact with the undergrowing skeletal muscle cells. After a week of co-culturing, we observed processes coming from motor neurons and establishing contact with underlying muscle cells (**Fig.5G** and **5H**).

This proof-of-principle demonstrates that VISIBLE effectively mitigates the inherent variability and complexity within co-culture system, thereby reducing experimental failure rates and enabling more complex investigations. Importantly, the diverse experiments presented here also showcase how real-time imaging-driven capabilities can be used to precisely guide bioprinting to significantly enhance the success and efficiency of experimental pipelines.

### Serial Intervention: scratch assay and recovery

Next, we set out to test the capability of our system to conduct a complete life-cycle of a complex experiment with serial intervention on the same sample with minimal human intervention. For this purpose, we culture and image skeletal muscle cells in VISIBLE before and after injury assessing the healing capabilities, a well-studied model in the field of wound repair and developmental biology ^38^.

First of all, we were able to carry out such complex experiment within the same machine and collecting the data on the fly, because VISIBLE has an incubation feature for culturing the induced scratch and recover within a week with minimal human intervention. Like this the plate is not moved or needs to leave VISIBLE. In this set of experiments, the manipulation tool-head was fixed with a nozzle (∼480 µm) and used to induce a controlled scratch in a well of confluent skeletal muscle cells (**Fig.6A**). A series of images were acquired in intervals of 24 hours by the microscope of the VISIBLE system demonstrating the recovery of the scratch within the system (**Fig.6B**). In the final step, differentiation medium was introduced via a syringe mounted on the tool-head. After 1 week, the final image demonstrates a fully recovered differentiated muscle cell culture following an induce scratch. The growth dynamics of the muscle cells during recovery phase (i.e., area vs time and gap width vs time) (**Fig.6C** and **D**) demonstrated a trend in line with previous studies performed with human intervention ^38, 39^. Furthermore, the entire procedure was performed by the VISIBLE system, hence eliminating the experimental variability of scratch size and positioning that usually is associated with these experiments when performed manually, and importantly, with this system we can guarantee the repeatability of the scratch assay across multiple timepoints, which would not be feasible by hand. We have also performed the scratch experiment simultaneously within 2 neighbouring wells to show the repeatability and high-throughput capabilities, also collecting live imaging data of the regrowth phase.

**Figure 6:**
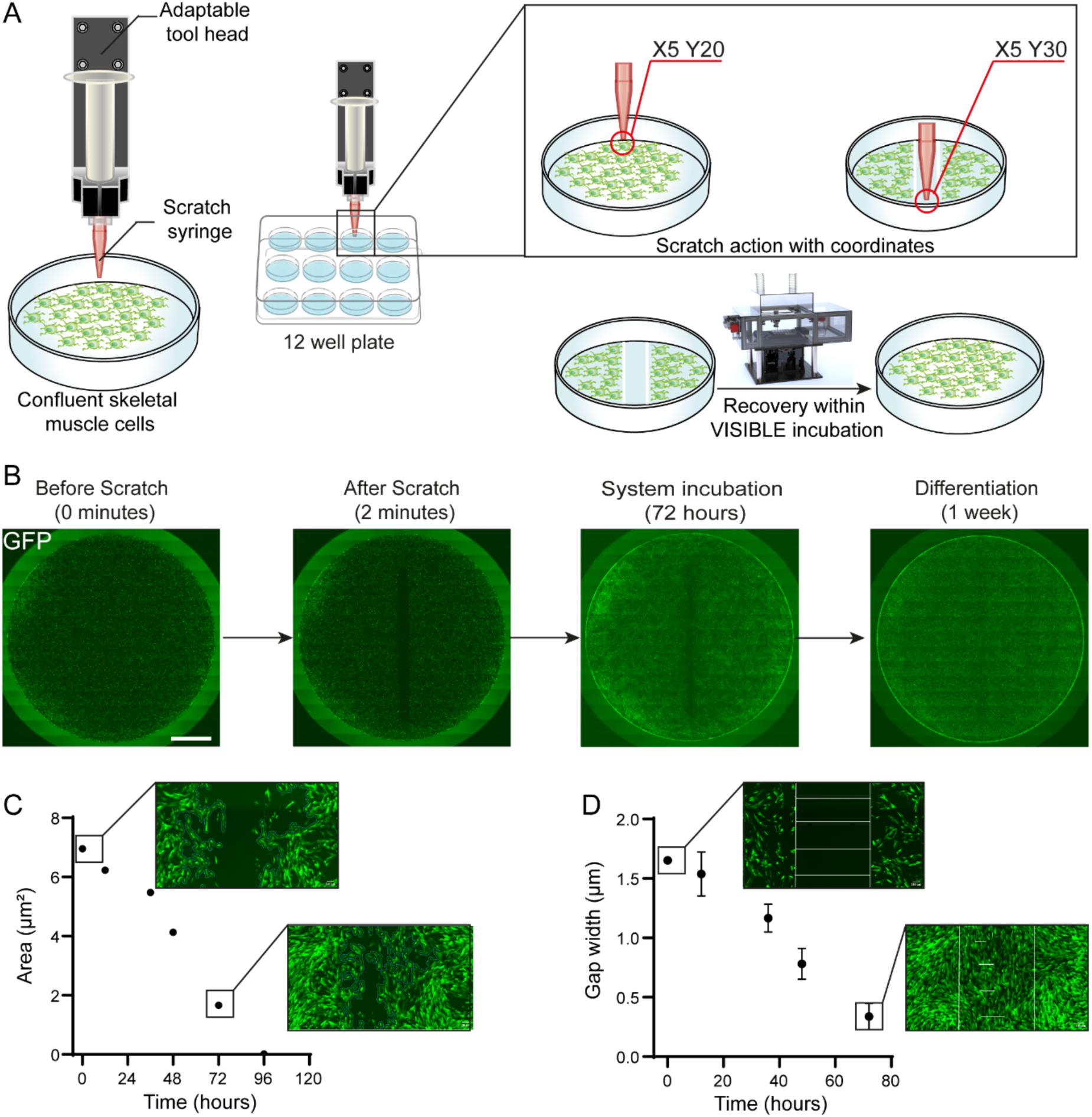
Serial intervention (scratch-assay on confluent muscle cells) (A) A schematic details the steps for performing a scratch assay experiment on confluent muscle cells inside the VISIBLE system. (B) An experiment demonstrating time-lapse progression of scratch (at 2 min) and healing of the scratch injury by >90% of scratch area recovery after 72 hours performed within the VISIBLE system, last image showcase differentiated muscle cells after 1 week of the initial scratch (Scale bar 1 mm). (C) Incubation ability of VISIBLE system as corroborated by area versus time graph, wherein within 72 hours of scratch injury muscle cells were proliferated and remarkably covered >90% of scratch area site. (D) Similarly, the gap width also demonstrated the same trend of scratch area recovery.

### pLarry barcoded organoid sampling in complex adherent cultures

We then wanted to test the ability of our system to integrate with complex preclinical translational pipelines which include multiple steps in vitro, ex vivo, and in vivo. For this, we have explored the feasibility of using VISIBLE system for patient-derived cancer organoids pipelines that require DNA barcoding of cells using lentiviral delivery enables lineage tracing and single-cell RNA sequencing to allow tracing of clones through phenotypic space^40^. However, optimizing conditions of a single barcode per cell may lead to incomplete infection of the cell population and consequently, negatively affect coverage of lineages within a population. Using a patient-derived xenograft organoid (PDXO) model of triple negative breast cancer^41^, we infected single cells with the pLARRY barcoding system and allowed organoids to form (**Fig.7A**). Infected organoids were detected by eGFP expression by the vision-based functioning of VISIBLE system and enriched (**Fig.7B**). To test whether organoids sampled through the VISIBLE system could be used for subsequent downstream analysis in vivo, we injected the sampled eGFP+ organoids into immune deficient mice. Tumours were collected and dissociated 5 weeks after injection (**Fig.7C**), followed by isolation of single cells and sequencing (**Fig.7D**). From tumours, we found 39% contained either eGFP+, the LARRY barcode or EEF1A promoter with barcodes detected in 1492 cells across 130 clonal identities (**Fig.7E**). Consistent with previous observations from this model^42^, we observed a subset of cells with high FGFR1 expression (**Fig.7E**). Intriguingly, we saw no difference in the Shannon diversity index between these subsets and the global population suggesting that phenotypic space is independent of clonal origin in the model of human triple negative breast cancer (**Fig.7F**). Altogether, this approach provides a key proof-of-concept to imaging and sampling subpopulations of cellular models that are viable for subsequent experimental use in vivo. Furthermore, it also shows how the VISIBLE system can be integrated in complex pipelines, enabling more precise experiments that also ultimately help to perform better in vivo experiments.

**Figure 7:**
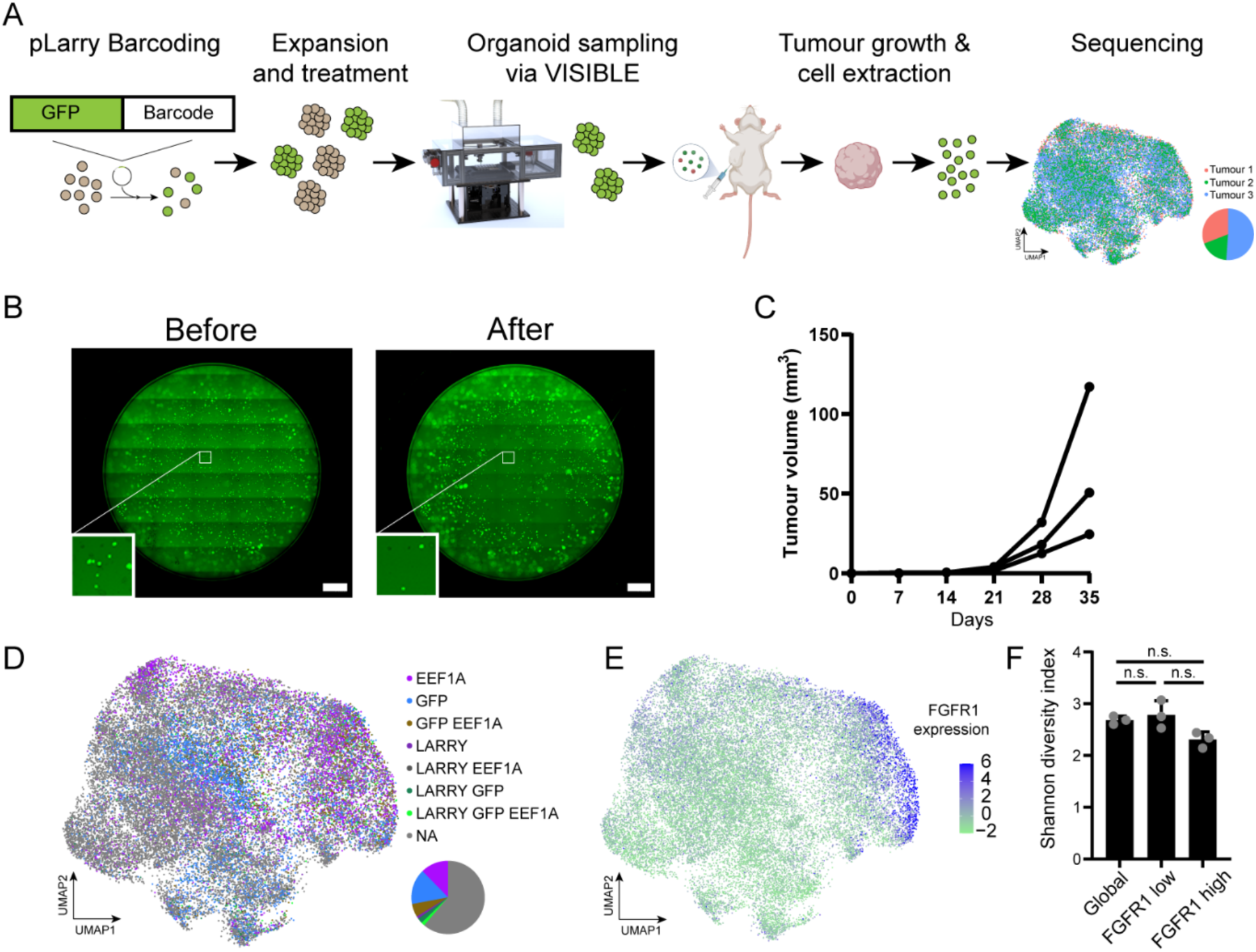
Enriching barcoded GFP+ organoids for tumour growth- (A) Schematic of experimental outline. (B) Images show wells of pLARRY GFP+ infected organoids before and after VISIBLE selection (Scale bar 1mm). (C) Volume quantification of tumours formed from selected organoids (n = 3 mice). (D) and (E). Uniform manifold approximation and projection (UMAP) representation of single-cell gene expression data generated from dissociated tumours. (D) Overlay indicates cells where elements of the pLARRY vector were detected. (E) Overlay indicates FGFR1 expression. (F) Bar graph depicting the Shannon diversity indices from the global, FGFR1 low (bottom 25% quartile) and FGFR high (top 25% quartile) populations.

## Discussion

In this study, we present a novel integrated, feedback-controlled platform that combines real-time imaging with automated manipulation and bioprinting to address the challenges of repeatability and heterogeneity in complex in vitro cultures. The VISIBLE system is capable of directly and iteratively manipulating live CIVMs cultures, and represents the ideal starting system for any lab interested in automating their culture or analysis for preclinical studies and basic research. Our system enables precise, iterative interventions guided by live image analysis, facilitating a wide range of experimental workflows from single-cell sampling to the handling of millimetre-scale hydrogels.

Throughout this paper, we have showcased the use of VISIBLE across a wide spectrum of different experimental set-ups, ranging from organoid sampling based on visible morphological differences (organoid diameters ∼ 1 mm), sampling of hiPSC derived neurospheres based on functional activity (calcium signalling), interactive 3D bioprinting, scratch assay on skeletal muscle cells and PDXO cancer organoids sampling for in-vivo implantation studies. Throughout all these different experimental examples, the use of VISIBLE allowed to directly interact with the cultures CIVMs in ways that would not have been possible with conventional culture systems or other platforms.

The modular architecture of the VISIBLE system also supports the tailoring of the platform towards bespoke applications and increases its versatility. For instance, the integration of multiple interchangeable tool heads enables scalable, high-throughput operations across entire multi-well plates with different commercially available attachments (i.e., anything from standard serological pipettes to mechanical probes and custom glass pipettes). The system is also designed specifically for live experiments, as it is enclosed in a custom incubator system with both temperature and environmental controls. Importantly, this aspect of the instrument is also customisable and allows for potentially hosting a number of different ancillary equipment such as, a multiplate holder for high-throughput studies.

Compared to other systems that allow for manipulation of complex cultures, our platform allows for simultaneous imaging and manipulation of live samples, which translates to its unique ability to perform imaging-guided sampling and bioprinting directly on live cultures. This has important implications, as it means that the manipulation, printing, sampling and generally any interaction with the cultured samples can be tailored on-the-fly and adapted to the dynamic condition of the culture, without transferring the plates across different instruments.

This unique set of features of the VISIBLE allows for example the precise selection of organoids from both suspension and adherent cultures, based on any feature or parameter that can be measured by imaging (e.g., morphology, expression of specific reporters, etc.). However, this also means that the inherent variability and dynamicity of CIVMs cultures is no longer a barrier, but can be effectively counteracted by adapting the different phases of the experiments to the status of the culture “as it is” rather than “as it is supposed to be”. One example of this is the experiment presented in Fig.4, where the variability in maturation speed of the human 3D neurosphere cultures (i.e., frequency of Ca^2+^ peaks variability between organoids within the same well) can be taken into account and the different maturation states “sampled” across different timepoints or separated in different collection samples.

We believe, this work substantially improves upon current experimental approaches for complex cultures, establishing a critical foundation for the development of “self-driving biological labs”. In fact, our system can be further improved by the planned future integration of machine-vision based “triggers” that could enable real-time event detection and autonomous decision-making, reducing human error and increasing experimental reproducibility.

Moreover, our current microfluidic-based suction mechanism—while capable of isolating single cells (∼50 µm)—can be further refined to minimise perturbation of neighbouring structures, improving precision in densely packed cultures such as hiPSC colonies.

The platform also supports advanced biological studies, such as serial scratch assays to investigate wound healing, bacterial biofilm formation, and other dynamic cellular processes with minimal manual intervention.

Ultimately, the VISIBLE system provides experimental biologists with enhanced control, continuous monitoring, and imaging-driven, iterative manipulation capabilities. Its modular, scalable design—compatible with widely available electronic and opto-mechanical components—offers unprecedented flexibility in customising in vitro experimental workflows. While this work highlights its application in neuroscience, 3D bioprinting, and cancer modelling, the VISIBLE system holds significant potential across a broad spectrum of disciplines, including drug screening, developmental biology, biofilm research, and biomechanics.

## Acknowledgments

**A.S.** wishes to acknowledge the support of the BBSRC (Grant BB/T011572/1 and Grant BB/W006561/1), and of the Dementia Research Institute (UKDRI). **F.S.T.** and **A.S.** acknowledge funding by the European Union (Horizon Europe project no. 101080690 – MAGIC). Views and opinions expressed are however those of the authors only and do not necessarily reflect those of the European Union or HADEA. Neither the European Union nor HADEA can be held responsible for them. This work is funded by the UK Research and Innovation (UKRI) under the UK government’s Horizon Europe funding guarantee grant no. 10080927, 10079726 and 10078461. The authors are grateful to the Myoline platform of the Institute of Myology (Paris, FR) for providing myoblasts. This research was funded in whole, or in part, by the Wellcome Trust. We would like to acknowledge the Making Lab facility, a Science Technology Platform at the Francis Crick Institute. **A.S.**, **A.I.**, and **F.S.T.** acknowledge support by the Francis Crick Institute, which receives its core funding from Cancer Research UK, the UK Medical Research Council (MRC) and the Wellcome Trust (CC0102). **E.S.** is supported by the Francis Crick Institute, which receives its core funding from Cancer Research UK (CC2040), the UK Medical Research Council (CC2040), and the Wellcome Trust (CC2040) and the European Research Council (ERC Advanced Grant CAN_ORGANISE, Grant agreement number 101019366). **E.S.** reports grants from Novartis, Merck Sharp Dohme, AstraZeneca and personal fees from Phenomic outside the submitted work.

**S.J.** acknowledges funding support from the BBSRC TRDF Grant (BB/T011572/1) and the Chris Banton Fund of The Francis Crick Institute, London. **S.J.** would like to thank the MedTech Super Connector (MTSC) fellowship and the BBSRC ICURe Explore Programme for funding support towards commercialization of innovation. **S.J.** is the recipient of LUSH Prize in the young researcher category and extends gratitude for generous funding support. **C.D.H.R.** is a recipient of a Bourse postdoctorale from the Fonds de recherche du Québec – Santé (https://doi.org/10.69777/273104), as well as an EACR-AstraZeneca Postdoctoral Fellowship. C.D.H.R. is supported by funding from the AstraZeneca-Crick Research Alliance.

The breast cancer patient-derived xenograft organoids PDXO GCRC1915 was obtained from the breast tissue and data bank (Park lab, McGill University) (PMID: 32546838), supported by the Réseau de Recherche sur le Cancer of the Fonds de Recherche du Québec-Santé and the Québec Breast Cancer Foundation, and certified by the Canadian Tumor Repository Network (CTRNet).

## Materials and Methods

### Assembling the VISIBLE system

The tool-head of manipulation system is mounted to an XY gantry whose motion is controlled by 2 stepper motors (Bipolar Hybrid Stepper Motor Nema 17) arranged in a core XY coordination. For the print-head displacements in Z-direction a customised in-house developed screw-based arrangement is employed (**Fig.1D**). Very precise movements of the printhead in the Z direction are vital, as even a minute displacement error in the downward Z-direction can potentially damage the objective lens of the integrated microscopic system. Moreover, as the present system is specifically designed to manipulate single-cell level precision, therefore Z-axis displacements are accurately controlled with end-stops/limit switches.

### Microscopic Unit

At the bottom sits the fluorescence microscope wherein objective lens directly see the glass slide/ tissue culture plastic plate placed on the printing bed. An automated stage with extremely precise stepper motors (C9004-9012K, Marzhauser) for allowing accurate XY positioning of the printing bed is controlled by an external high-resolution positioning controller (Corvus, ITK). This system comes with its joystick controller for stage positioning, but we have interfaced the stage controller (drivers and .dll files for Corvus) with an open-source Micromanager allowing automation-based control of our system. Therein lies a filter cube (ThorLabs) with excitation and emission filters for DAPI/FITC/Texas Red (Semrock). The filter set is illuminated by a light source (CoolLED, pE-300) wherein light is transmitted through liquid light guide, collimating adapter (ThorLabs LLG3A5-A), and lens arrangement. Finally, a CMOS compact scientific camera (Kiralux 12.3 MP) sits at the opposite end to capture images. To control and coordinate simultaneous functionality of different fluorescence microscope components, we have interfaced these components with Micro-Manager software (version 2.0 for 64-bit system). Suitable software drivers and .dll files for stage controller, CoolLED light source, and CMOS camera were loaded into the system for their respective operations. As a result, we have developed an open-source system wherein the control operations can be customized based on the end-user’s specific experimental requirements.

### The Integration Hardware

At the heart of the system sits the central controlling unit (Smothieboard V2), which is 32-bit open-source firmware comprising of 5 stepper drivers with 1/32 micro stepping capability. This controlling unit is powered by a 24 V power supply encased in a polycarbonate enclosure and is responsible for controlling the stepper motors and limit switches of the bioprinter unit. The controlling unit is interfaced with the PC through an Ethernet cable and the commands for XY movement of the print-head are encoded in the G-code. Additionally, other programming languages, for instance MATLAB or Python can be employed to send commands to the 3D printing unit.

The entire gantry system for 3D bioplotter is mounted on the squared solid aluminium optical breadboard (matte black anodised finish, ThorLabs) with dimensions of 45 cms x 45 cms. At the centre of this breadboard, a rectangular opening of dimensions of 18 cms x 14 cms was custom-cut for arranging the objective lens and associated set-up for the fluorescence microscope. The bottom microscope assembly is also assembled on a squared solid aluminium optical breadboard (matte black anodised finish) with dimensions of 45 cms x 45 cms. There are 4 pillar posts with M4 tapped holes on both ends (Pedestal Pillar Posts, Thor Labs) to support the top 3D bioprinter breadboard (height 32 cms). We call this method as imaging-driven manipulation. The system uses a top object-identifier/locator camera for well plate identification and a bottom inverted epifluorescence microscope for single-cell level resolution.

To maintain the temperature requirement, regulated hot air is continuously supplied via a controlled chamber The Cube 2 (Life Imaging Services). A metallic sensing element based on Resistance Temperature Detection (RTD) is placed at the height of 3D printing stage for indicating the accurate temperature in the vicinity of the cell environment. The overall dimension of the 3D bioprinting chamber along with the top polycarbonate-based enclosure is 52 x 45 x 32 cms.

### Skeletal muscle cell culture and analysis

#### Myoblast cultures

Human immortalised skeletal myoblasts (AB1167) were kindly provided by the Myoline platform of the Institute of Myology in Paris, France. Cells were transfected with a lentivirus with a GFP reporter to facilitate live visualisation and expanded in skeletal muscle cell growth medium (Promocell) at 37 °C, 5% CO2. For co-culture and scratch experiments, cells were seeded at a density of 21000 cells/cm^2^ in Matrigel (Corning)-coated plates, once reached the appropriate confluency, growth medium was changed into differentiation medium composed of DMEM high glucose (Sigma), supplemented with 1% Glutamax (Thermo), 1% pen/strep (Thermo) and 10 µg/ml insulin (Gibco). After 4 days, mCherry NSs were deposited on highly differentiated myoblast areas, in parallel, skeletal muscle differentiation medium was diluted in 50/50 medium at a 1:1 ratio. The co-culture was maintained until day 7.

#### Analysis of differentiation areas and NSs coverage

Acquired images were processed in Fiji, where 5 regions of interest (ROIs) were considered for each photo and manually distributed in low- and high-density myogenic areas. For each ROIs, a selection was generated to quantify the areas covered by the myogenic cells. The data were then expressed as a percentage of the total ROI area. For co-culture experiments, the areas covered by the NSs were calculated following the same process.

#### Scratch test

To perform scratch tests with VISIBLE system, a 12-well plate with 80% confluent skeletal muscle cells was kept in the system. A syringe with a diameter of (∼150 µm) was attached to the print-head to introduce the scratch. First, the coordinates of the start point (X5, Y20) were recorded and then to introduce a straight scratch end point coordinates (X5, Y30) were executed via the system. This process generated a clean straight scratch injury in the expanding muscle cell culture (**Fig.6B**). This process of introducing a scratch injury took 2 minutes and then the plate was incubated in the system and imaged at an interval of 6 hours until full recovery.

#### Scratch test analysis

Live captured images were analysed in Fiji. Cells’ ability to recover from injury was measured as the area and gap width changing over time. Selections were generated to quantify the total empty space or the gap width at the different time points.

#### Neurosphere generation and culture

Human iPSC (hiPSC)-derived NGN2-induced neurons were generated using a doxycycline inducible system from piggyBac-mediated stable integrated NGN2 iPSCs (cell line: KOLF2.1J). Neurospheres were generated as previously described with modifications^34^. Single hiPSCs were plated on a PDMS-based microwell device with well sizes 400 µm width x 400 µm length, in a 24-well plate at 1.4 x 10^6^ cells per well. The cells were maintained in induction media consisting of KnockOut DMEM/F12 (Thermo), N2 supplement (Thermo), NEAA (Thermo), mouse laminin (1 µg/ml, Thermo), ROCK inhibitor (10µM, Tocris) and doxycycline (2 µg/ml, Sigma) at 37 °C, 5% CO2. After 24 hours, the ROCK inhibitor was removed from the culture. Neurospheres were then harvested and transferred to a10-mm dish with orbital shaking at 60 rpm. On day 3 of induction, the medium was switched to maturation media, consisting of 50% Neurobasal (Thermo) with B27 supplement (Thermo) and 50% Advanced DMEM/F12 (Thermo) with N2 supplement, along with Glutamax (Thermo) and Pen/Strep (Thermo). The neurospheres were plated on a Matrigel (Corning)-coated plate and allowed for maturation for 16 days before functional-based selection experiments were conducted.

#### Neuronal activity analysis

The plated neurospheres were incubated with the Ca2+ dye Fluo-4AM (5 µM, Thermo) for 1 hour at 37 °C, 5% CO2, subsequently washed with PBS and replaced with maturation medium. For analysis, calcium images were processed using ImageJ software for bleach correction, ROI selection, and intensity measurement. After correcting bleaching, five circular ROIs with a 3-pixel radius were drawn on the cell soma in each neurosphere to measure mean intensity across the recorded images. The intensity data were then input to the peakcaller^43^ to acquire the number of calcium peaks. Representative calcium traces for each ROI were calculated using the formula dF/F = (F-F0)/F0, where F is the current fluorescence value and F0 is the mean fluorescence value of the first 5 frames.

#### Hydrogel fabrication

For constituting hydrogels, all weighing measurements and mixing procedural steps were performed in sterile conditions inside a bio-safety cabinet. Glass beakers and stirring magnets were autoclaved. Preparation steps started by measuring neuronal culture medium (4 mL) and homogenously mixing it with Matrigel (1 mL). Hyaluronic acid (5 mg/mL) from Sigma Aldrich was mixed in the above solvent and stirred overnight (12 hours) at ambient temperature. This was followed by the addition of Fibrinogen (45 mg/mL) from Sigma Aldrich and stirred for 5 hours at room temperature. As a final step, Alginate (5% (w/v)) from Sigma Aldrich is added to the above mixture and stirred overnight to obtain a homogenous hydrogel matrix. Freshly prepared hydrogel was used in our present experimental studies, but as-prepared hydrogel matrix can be stored at -20 °C for 6 months.

Motor Neuron Progenitors cells (MNPs) were dissociated from a 6-well plate using mix of EDTA in PBS and were gently mixed with the as-prepared hydrogel matrix (with concentration of 4-5 Million cells/mL) by pipetting them up-down gently for homogenous distribution of cells throughout the hydrogel, leading to the formation of a bio-ink. This bio-ink was crosslinked by a 50:50 solution consisting of calcium chloride (CaCl2) (1.5% (w/v)) and thrombin (25 U/mL in 0.1% BSA Solution). Bio-ink was crosslinked for 15 minutes at room temperature. After crosslinking, the bio-ink was washed with PBS for 3 times. This was followed by flooding the wells gently through the walls of a cell plate with neuronal culture media supplemented with Compound E and kept in an incubator at 37 °C with 5% CO2.

#### LARRY lineage tracing

Detailed patient-derived breast cancer organoid culturing conditions and experimental methods can be found in Ratcliffe et al, BioRXiv 2025. Key steps are summarised below. GCRC1915 PDXO parental organoid lines were generated from the GCRC1915Tc PDX^41^ and authenticated, as well as tested for mycoplasma contamination by the Cell Services Platform at the Francis Crick Institute. LARRY Barcode Version 1 library was a gift from Fernando Camargo (Addgene #140024) and was used to prepare lentiviral particles. GCRC1915 PDXO cells were infected with an MOI of 50 in 200 µL culture media lacking Cultrex under culture conditions (37°C, 5% CO2, humidified atmosphere) for 1 hour. Cells were then split into 3 and plated. After 7 days of culture, organoids were picked using VISIBLE.

#### In vivo experiments

The Francis Crick Institute’s Animal Welfare and Ethical Review Body and UK Home Office authority provided by Project License 0736231 approved all animal model procedures. Procedures described in this study were compliant with relevant ethical regulations regarding animal research. Using the VISIBLE system, GFP+ organoids were picked, spun down and resuspended in Matrigel (Corning Cat No. 354234) prior to injection. After 5 weeks, tumours were collected, dissociated and human tumour cells were enriched using a Miltenyi tumour dissociator according manufacturer’s instructions.

#### Single cell processing, sequencing and analysis

Cell concentration and viability was measured and approximately 120,000 cells per tumour were loaded on a Chromium Chip and processed according to manufacturer’s instructions (CG000315 Chromium Single Cell 3’ Reagent Kits User Guide (v3.1 - Dual Index)) to generate cDNA libraries using Chromium Next GEM Single Cell library reagents Final libraries are QC’d using the Agilent TapeStation and sequenced using the Illumina NovaSeq 6000. Sequencing read configuration: 28-10-10-90.

The GFP-LARRY plasmid was added to the reference genome refdata-gex-GRCh38-2020-A and the GTF annotation of the plasmid transcripts was added to the same reference annotation file. Cellranger (version 7.0.1) mkref was run to create a new reference for genome annotation. Cellranger count was used to count the 10x libraries. A whitelist of cell barcodes was obtained with UMI-tools (version 1.1.2)^44^ and the cellranger quantifications were imported into a Seurat (version 4.3.0)^45^ object. Doublets were calculated using scDblFinder (version 1.8.0)^46^ and the LARRY barcode information was added to the Seurat objects. The R programming language was used (version 4.1.2) (R Development Core Team 2008). We investigated key QC parameters, and we removed cells with a low number of detected features or cells with a very high proportion of mitochondrial gene expression. We selected a lower bound filter for both the minimum number of reads per cell and the minimum number of detected features using 3 median absolute deviations (MADs). For the percentage of mitochondrial genes, we set upper bounds using 3 MADs. We use the “SCTransform” method (version 0.3.5)^47^ for the normalisation and variance stabilisation. The dataset was subsetted according to the lowest 25% and highest 25% FGFR1 expressing cells. In the global population, as well as within subsets, barcodes were used to determine the Shannon diversity index according to:

H = - ∑ (pi * ln(pi))

Where H is the Shannon diversity index, pi is the proportion of a clone belonging to the barcoded population.

