## Supplemental Info for "VISIBLE: An imaging-driven system for sampling, biofabrication, and manipulation of complex biological models" (from another version of this preprint: version 1, 2025-06-17)

### Supplementary Information:

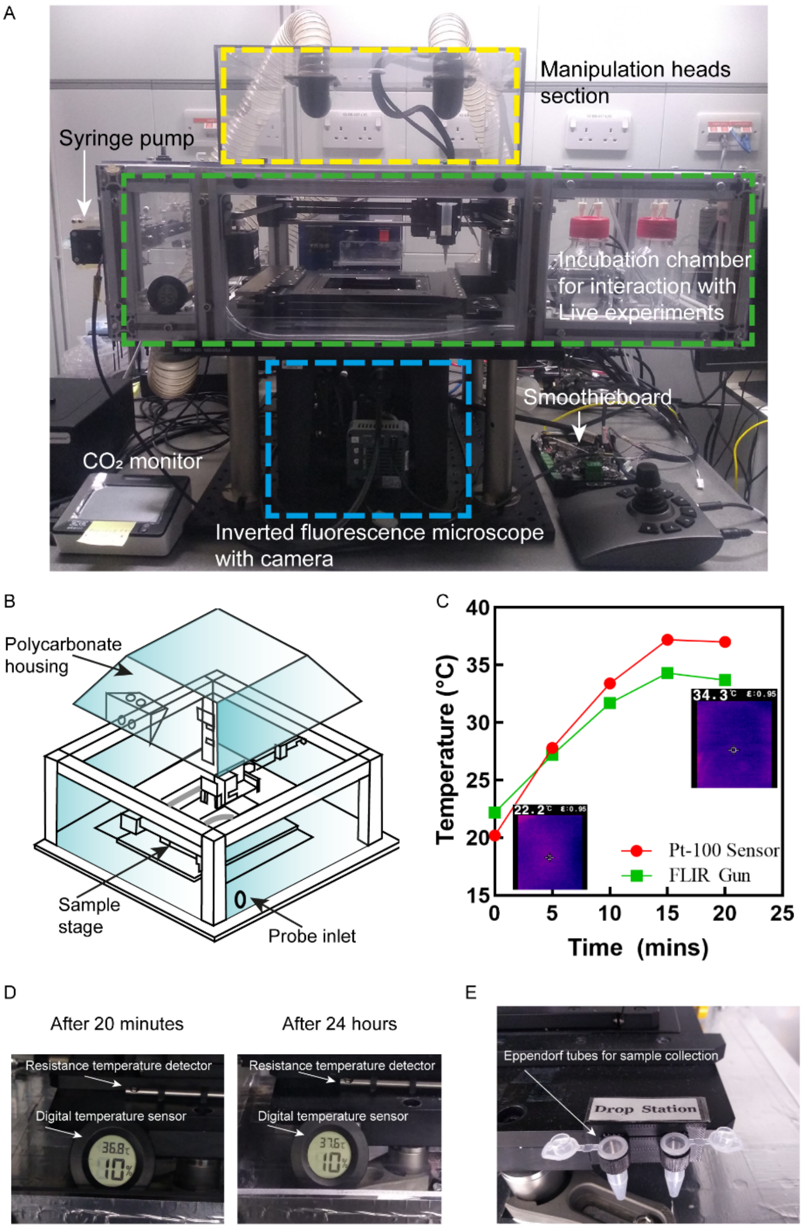


***Supplementary Figure 1: Engineering the VISIBLE system****– (A) A photograph of the VISIBLE system with its different components and their respective functions. (B and C) VISIBLE system is housed in a transparent polycarbonate housing for ensuring sterile conditions for complex biological experiments and temperature characteristics (37 ℃ and 5% $CO_2$) suitable for performing in vitro studies measured by Pt-100 sensor (Resistance Temperature Detector, RTD) and Forward Looking Infrared (FLIR) gun. (D) Digital temperature sensor (mini temperature LCD monitor) placed inside the VISIBLE system showcases the temperature retaining capability of VISIBLE system for 24 hours. (E) Eppendorf tubes stationed on the stage for biological sample collection for downstream analysis.*

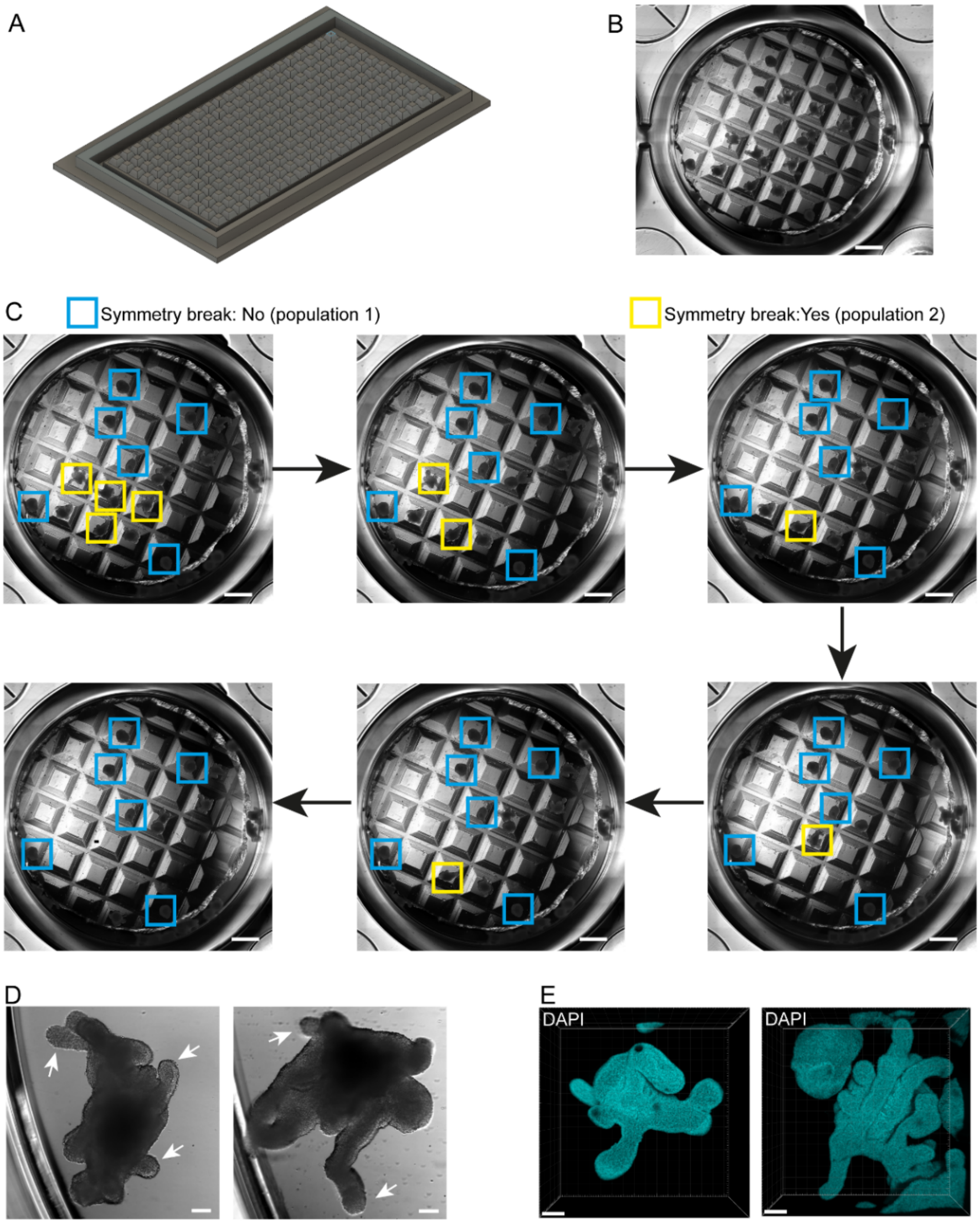


***Supplementary Figure 2: Morphology-based sampling of suspended neuroectodermal organoids***- *(A) A 3D printed cast of custom wells designed for realising a complex heterogenous culture of organoids in suspension. This cast was specifically designed and developed for conducting sampling experiments via VISIBLE system. (B) A stitched image generated by VISIBLE system showcasing an example of a complex heterogenous organoid culture, wherein organoids with and without broken symmetry are randomly distributed in PDMS wells (Scale bar 1 mm). (C) A sequential iterative sampling of organoids that have broken symmetry (population 2) were collected by the VISIBLE system. (D) Microscopic images show picked and transferred organoids on a collection plate. Arrows (white) indicate intricate details of organoids that remains preserved during the sampling and collection process (Scale bar 200 µm), and (E) Downstream analysis (ICC studies) were performed on the collected organoids indicating the suitability of VISIBLE system for sampling experiments in complex in vitro cultures (Scale bar 150 µm).*

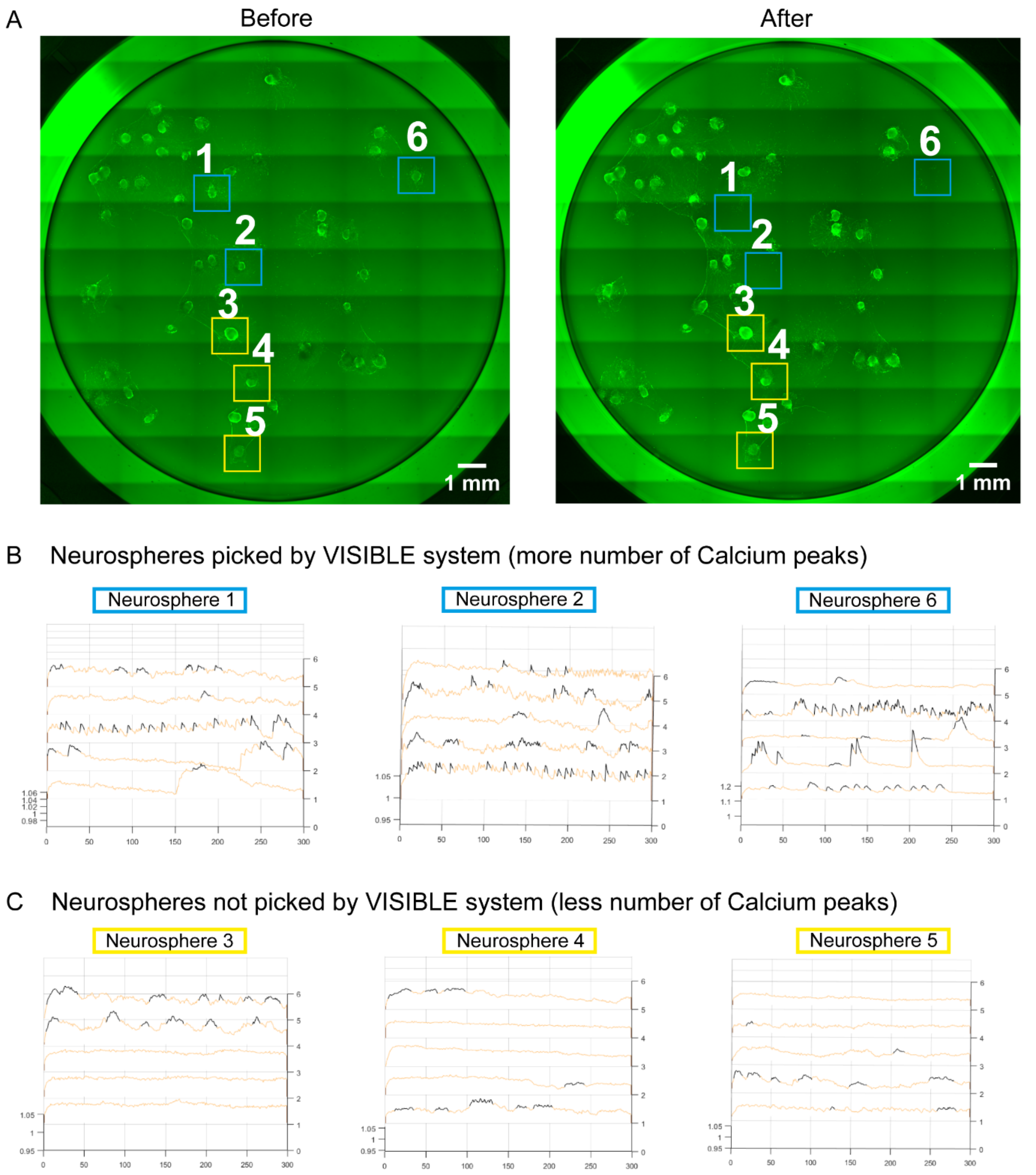


***Supplementary Figure 3: Function-based sampling of adherent neurospheres in complex culture**- (A and B) Before and after stitched image of the experimental well wherein, a complex culture of neurospheres is used for sampling based on calcium signalling (Scale bar 1mm). (B and C) Graphs indicating number of Calcium peaks from 6 different ROIs from a single neurosphere. (B) Neurosphere numbered (1, 2 and 6) showed more numbers of Calcium peaks and were sampled by the VISIBLE system. (C) Neurosphere numbered (3, 4 and 5) showed a very few numbers of Calcium peaks and these remained in the culture unperturbed by the VISIBLE system.*

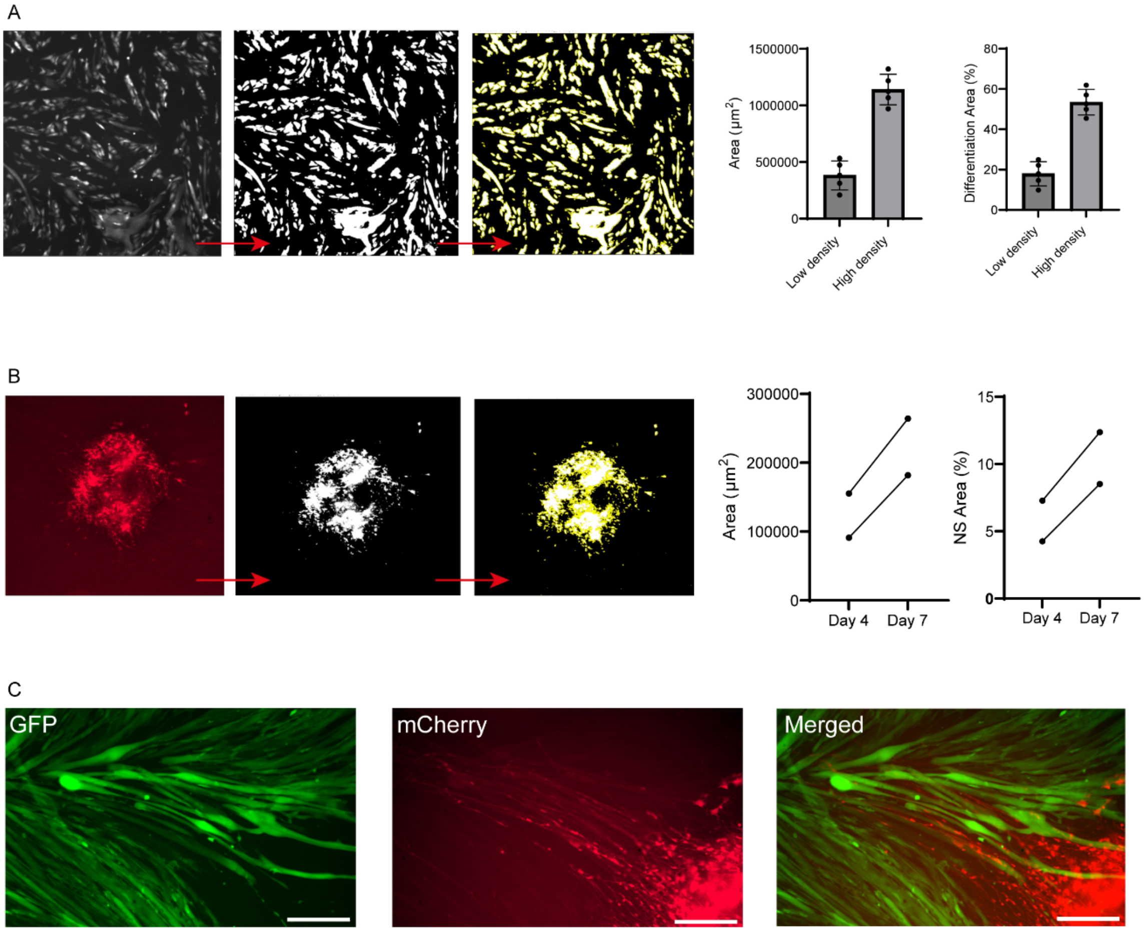


***Supplementary Figure 4: Serial intervention for spatiotemporal experiments-*** *(A) Analysis of coverage area of skeletal muscle cells to identify areas of low and high density for deciding of location to drop neurospheres for spatiotemporal experiments. (B) Analysis of coverage area by neurosphere and its processes over skeletal muscle cells after day 4 in culture. (C) A higher magnification image of co-culture wherein effective interaction between dropped neurosphere and underlying skeletal muscle cells is observed after 7 days in culture (Scale bar 200 μm).*

**Supplementary Videos 1 to 3 are provided in FigShare:**

**https://doi.org/10.6084/m9.figshare.29310578.v1**
